# Discriminating models of trait evolution

**DOI:** 10.1101/2025.06.12.659377

**Authors:** Jenniffer Roa Lozano, Michael DeGiorgio, Raquel Assis, Rich Adams

**Affiliations:** Center for Agricultural Data Analytics, University of Arkansas, Fayetteville, AR; Department of Entomology and Plant Pathology, University of Arkansas, Fayetteville, AR; Department of Electrical Engineering and Computer Science, Florida Atlantic University, Boca Raton, FL; Institute for Human Health and Disease Intervention, Florida Atlantic University, Boca Raton, FL

**Keywords:** Comparative biology, phylogenetics, trait evolution, gene expression, comparative genomics, machine learning

## Abstract

A central challenge in comparative biology is linking present-day trait variation across species with unobserved evolutionary processes that occurred in the past. In this endeavor, phylogenetic comparative methods are invaluable for fitting, comparing, and selecting evolutionary models of varying complexity and biological meaning. Traditionally, evolutionary studies have relied on conventional statistical approaches to assess model fit and identify the one that best explains variation in a given trait. Here we explore an alternative strategy by applying supervised learning to predict evolutionary models via discriminant analysis. We formally introduce Evolutionary Discriminant Analysis (EvoDA) as an addition to the biologist’s toolkit, offering a suite of new methods for studying trait evolution. We evaluate the performance of EvoDA alongside conventional model selection through a series of fungal phylogeny case studies, each targeting increasingly challenging analytical tasks. These results showcase the strengths of EvoDA, with substantial improvements over conventional approaches when studying traits subject to measurement error, which likely reflect realistic conditions in empirical datasets. To complement our simulation-based benchmarking, we explore the application of EvoDA for tackling a notoriously difficult task: predicting the mode and tempo of gene expression evolution. This empirical analysis suggests that stabilizing selection acts on a majority of genes, with bursts of expression evolution in a handful of genes related to stress, cellular transportation, and transcription regulation. Collectively, our findings illustrate the promise of EvoDA for predicting trait models across a range of evolutionary and experimental contexts, establishing a new methodological framework for the next era of comparative research.

**Significance Statement:** To make sense of biodiversity, evolutionary studies have historically relied on conventional statistical procedures to evaluate competing hypotheses about the mode and tempo of trait evolution. Here, we introduce new supervised learning methods that substantially outperform traditional techniques for correctly assigning trait models across a range of evolutionary and experimental conditions. We find that these methods are highly robust to measurement noise expected from realistic trait data and offer new insights into a central question in comparative genomics: what are the evolutionary forces shaping variation in gene expression across species?

## Introduction

Phylogenetic comparative methods (PCMs) have been a defining force in shaping modern evolutionary biology. Over the past few decades, they have ignited a renaissance in the study of large-scale biodiversity patterns and the processes driving them (Blomberg and Garland 2002; Pennell and Harmon 2013; Cornwell and Nakagawa 2017). At the core of PCMs are statistical models that define the probability distribution of trait changes along the branches of a phylogeny, governed by a set of parameters designed to capture key processes influencing trait evolution over time (Butler and King 2004; Beaulieu et al. 2012; Uyeda et al. 2018). Many of these methods are grounded in the foundational Brownian motion (BM) model and its extensions (Felsenstein 1985; Martins and Hansen 1997), most notably the Ornstein-Uhlenbeck (OU; Hansen 1997) and Early-Burst (EB; Harmon et al. 2010) models. These frameworks have been expanded to incorporate shifts in ancestral traits (Blomberg et al. 2020; Adams et al. 2021), multiple optima (Pennell et al. 2014), and additional features of evolution and diversification (Adams and Collyer 2018; Adams et al. 2019; Dimayacyac et al. 2023).

Because PCMs require an explicit model of trait evolution, a primary goal of comparative studies is to identify the model that best explains evolutionary variation in a studied trait. As with any model-based inference, selecting an appropriate model is typically considered a critical first step toward accurate inference (Felsenstein 1985; Blomberg and Garland 2002; Pennell and Harmon 2013; Uyeda et al. 2018). Importantly, when an assumed model contains too few parameters, important processes may be overlooked (Sullivan and Swofford 2001; Bengtsson and Cavanaugh 2006; Silvestro et al. 2015). Conversely, when a model is overly complex with too many parameters, inferences can also be unreliable (Huelsenbeck et al. 2001; Boettiger et al. 2012; Uyeda et al. 2018). The inherently model-based perspective of PCMs provides a natural framework for balancing this bias-variance trade-off by weighing the evidence for a model in relation to its complexity (Huelsenbeck et al. 2001; Uyeda et al. 2018).

There has long been a predominant focus on conventional model comparison and selection procedures for PCMs, particularly those based on information criteria applied using maximum likelihood (Goloboff and Arias 2019) or Bayesian inference (Holder and Lewis 2003; Gelman et al. 2014). Consequently, a multitude of strategies have emerged for identifying the most suitable evolutionary model from a set of plausible candidates. A widely used approach involves comparing fitted models based on their likelihood scores, penalized by the number of parameters to mitigate overfitting (Goloboff and Arias 2019). Prominent examples include the Akaike information criterion (AIC; Akaike 1973), Bayesian information criterion (BIC; Schwarz 1978), and corrected AIC (AICc; Hurvich and Tsai 1989), which are widely regarded as standard in the field (Butler and King 2004; Posada and Buckley 2004). These methods seek to strike a balance between complexity and goodness-of-fit (Posada and Buckley 2004; Jhwueng et al. 2014), with the total number of parameters playing a key role when comparing models of differing biological applications and interpretation. This issue is particularly relevant for popular PCM models that differ in likelihood function but share the same number of parameters, such as the stationary Ornstein-Uhlenbeck (Hansen 1997; Tung Ho and Ané 2014), Early-Burst (Harmon et al. 2010), and Pagel’s Kappa, Lambda, and Delta models (Pagel 1999). Additionally, several Bayesian strategies have been employed for selecting trait models (Huelsenbeck et al. 2001). These approaches benefit from their ability to naturally accommodate uncertainty in the posterior distribution and enable calculation of marginal likelihoods (Huelsenbeck et al. 2001; Susko and Roger 2020), though they are also notoriously demanding in time and computational investment.

Alternative strategies for classification tasks, which have largely remained unexplored in comparative studies, have gained significant traction in other areas of biology (Tarca et al. 2007; Okser et al. 2014; Libbrecht and Noble 2015). In particular, supervised learning has become transformative for achieving high-accuracy prediction and inference across many fields and disciplines (Cunningham et al. 2008; Zhou 2018; Żurański et al. 2021). Recent applications in evolutionary biology include phylogenetics (Mo et al. 2024; Silvestro et al. 2024), population genomics (Schrider and Kern 2018; Korfmann et al. 2023), functional genomics (Rives et al. 2019; Washburn et al. 2019; Routhier and Mozziconacci 2022), and systems biology (Bacardit et al. 2009). Contrasting with conventional model fitting, supervised learning seeks to build a predictive function *ŷ* = *f*(**x**), where the output response variable *ŷ* is predicted from a vector of input feature variables **x**. By utilizing a set of labeled training data with known **y** and **x**, supervised learning optimizes *f*(**x**) to minimize the discrepancy between predicted outcomes *ŷ* and true labels **y**. This emphasis on minimizing prediction error contrasts with the more conventional focus of PCM model fitting and comparison.

In this study, we explore the application of supervised learning to predict models of trait evolution using discriminant analysis (Fig. 1). Discriminant analysis encompasses a family of supervised algorithms that are trained to distinguish among classes by learning the boundaries that separate them (Larrañaga et al. 2006). It has been successfully applied in many fields for pattern recognition (McLachlan 1992) and dimensionality reduction (Ye and Ji 2009). Here, we investigate the potential of discriminant functions as a new strategy for predicting the evolutionary models underlying trait variation across species. We formally introduce Evolutionary Discriminant Analysis (EvoDA) as an addition to the PCM toolkit and apply it to a series of fungal phylogenetic case studies to evaluate its predictive performance under a range of evolutionary and experimental conditions. A key focus is the assessment of model selection accuracy for both EvoDA and conventional AIC-based strategies when analyzing noisy trait data subject to measurement error (Susko and Roger 2020; Bartoszek et al. 2023). That is, we assess how well EvoDA predicts evolutionary models when traits have been measured imprecisely, an issue that is highly relevant in empirical studies (Hansen and Bartoszek 2012; Silvestro et al. 2015). Complementing this work, we apply EvoDA to an empirical case study to predict mechanisms of gene expression evolution in the same fungal system—a task that has proven difficult for conventional PCMs (Cope et al. 2020; Dimayacyac et al. 2023). Collectively, we seek to understand both the promise and pitfalls of discriminant functions for evolutionary model selection.

**FIGURE 1.**
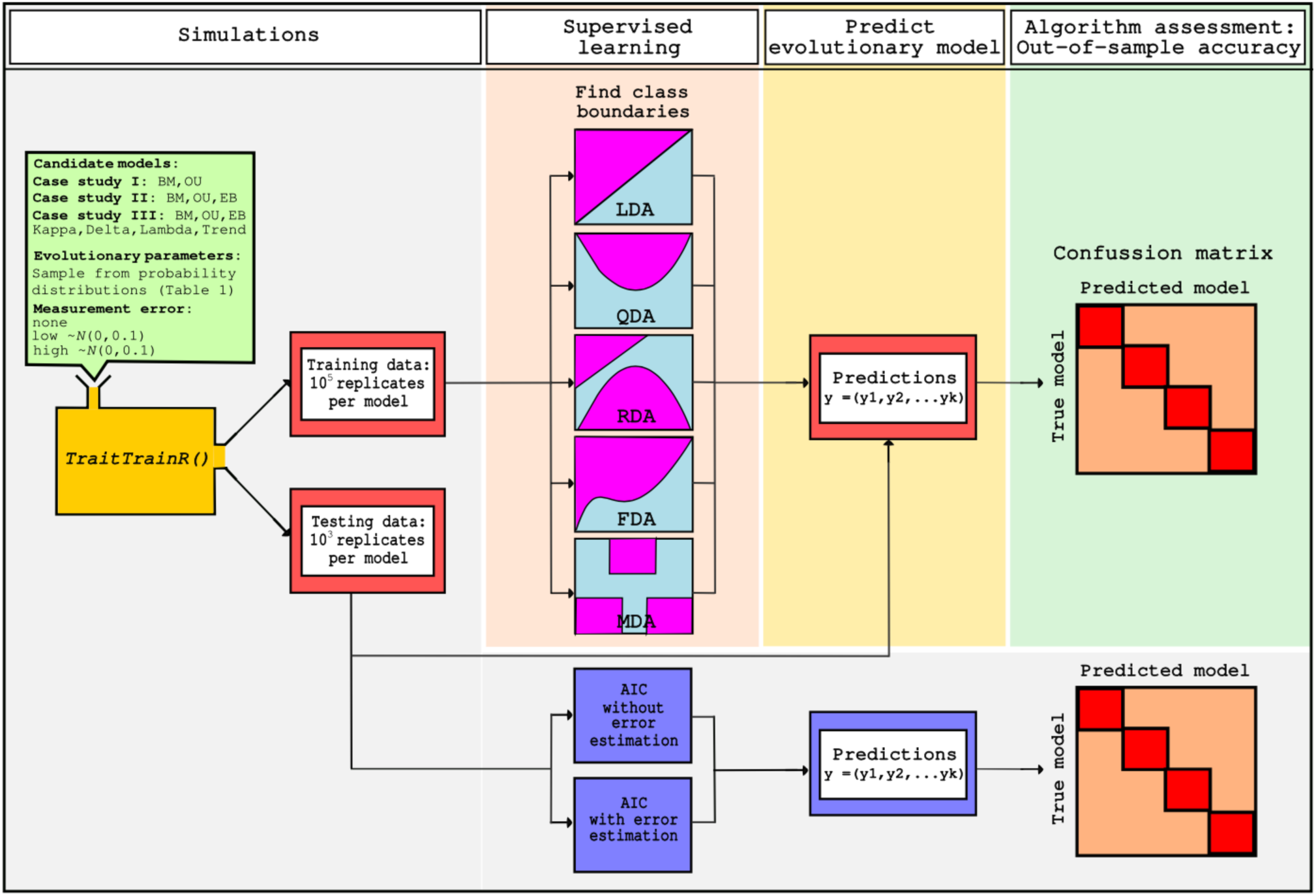
Workflow for evolutionary model classification using supervised learning with EvoDA, alongside conventional AIC-based model selection with and without error estimation. Simulated training and test datasets were generated using TraitTrainR under specified sets of candidate evolutionary models and parameter values.

## Results

### Study design: exploring EvoDA performance across three increasingly difficult case studies

We explored the performance of EvoDA in three phylogenetic case studies of escalating difficulty to evaluate model selection accuracy under varying evolutionary and experimental conditions. Five EvoDA algorithms were implemented and evaluated (see *Methods*): linear discriminant analysis (LDA; Sudibyo et al. 2020; Zaki and Meira, Jr 2020), quadratic discriminant analysis (QDA; Finch and Schneider 2006; Sifaou et al. 2020; Ghosh et al. 2021), regularized discriminant analysis (RDA; Zhu and Huang 2013; Yan et al. 2022), mixture discriminant analysis (MDA; Reynès et al. 2006), and flexible discriminant analysis (FDA; Ramsay and Silverman 2005). In each case study, we compared EvoDA accuracy alongside conventional model selection using two variants of AIC that differed only in whether measurement error was estimated or not (“AIC with measurement error” and “AIC without measurement error”). We focused our three simulation case studies using a fungal phylogeny of 18 species spanning over 800 million years (MY) of divergence (Cope et al. 2020), which has previously served as a benchmark for evaluating evolutionary model selection strategies (Cope et al. 2020) and investigating gene expression evolution (Dimayacyac et al. 2023).

The three case studies were structured as increasingly difficult classification tasks with either two (Case Study I), three (Case Study II), or seven (Case Study III) candidate models. Each case study reflects a distinct set of questions that are commonly addressed in comparative analyses. For example, a popular goal is to determine whether a trait evolved via BM or OU processes (Tung Ho and Ané 2014; Jhwueng and Maroulas 2016), reflecting Case Study I. In Case Study II, we sought to discriminate among the BM, OU, and EB models, which is a common question for gene expression data, for example (Dimayacyac et al. 2023). Finally, Case Study III tackled a more challenging analytical task spanning all seven canonical models (BM, OU, EB, Kappa, Lambda, Delta, and Trend). Measurement error is a ubiquitous issue in modern trait studies, and even small amounts are known to compromise model performance in many PCMs (Ives et al. 2007; Silvestro et al. 2015). For each case study, we therefore conducted three additional sets of analyses incorporating increasing amounts of measurement error in the trait data, which was accomplished by sampling noise from a normal distribution with mean zero and a standard deviation of 0.1 (“low error”) or 1.0 (“high error”), in addition to the “no error” scenario. Across all three case studies and associated analyses, RDA consistently converged to QDA with proportion *α* = 1 during hyperparameter tuning, yielding identical predictions and accuracy for both algorithms. Accordingly, we report and discuss their results jointly as QDA/RDA. Collectively, our three case studies each compared the performance of three error settings (“no error”, “low error” and “high error”) for each of seven model selection techniques (AIC with error estimation, AIC without error estimation, LDA, QDA, RDA, MDA, and FDA).

### Case Study I: discriminating BM and OU

Conventional model selection with AIC showed variable performance, with accuracy ranging from 43.9 to 87.0% across conditions and error settings (Fig. 2). Performance differed markedly between BM and OU models, depending on the application of error estimation and presence of measurement error in the trait data. Without error estimation, AIC achieved high accuracy for OU (99.7%) but performed poorly for BM (67.0-74.3%). Conversely, with error estimation, accuracy for BM improved (79.4-80.9%) but at a dramatic cost of substantially lower accuracy for OU (7.7 to 8.5%). Overall, AIC was highly sensitive to measurement error, with reductions in accuracy largely stemming from its inability to consistently predict BM without error estimation or OU with error estimation.

**FIGURE 2.**
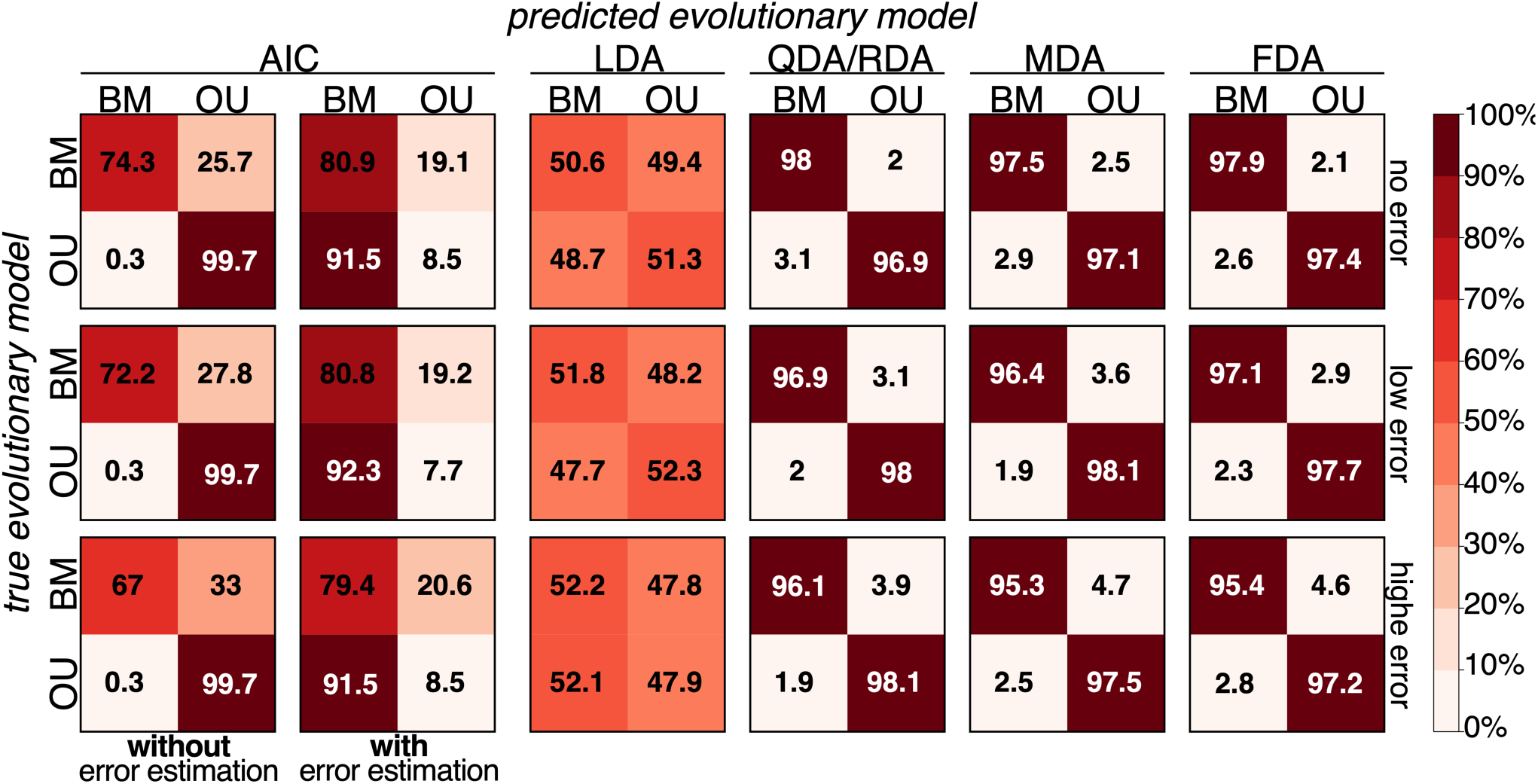
Case Study I: Discriminating between two canonical models of trait evolution. Confusion matrices (left to right) depict classification accuracies for AIC without and with error estimation, LDA, QDA, RDA, MDA, and FDA. QDA and RDA produced identical results and were therefore combined as QDA/RDA. Results are shown for experimental conditions with three levels of simulated trait measurement error: none (top row), low (middle row), and high (bottom row). In each matrix, rows represent the true evolutionary model used to simulate traits, and columns represent the predicted model. Shading intensity reflects the percentage of test replicates predicted for each model, with darker cells indicating higher values.

In contrast, all five EvoDA algorithms exhibited consistent performance across conditions and error settings (Fig. 2). LDA performed poorly, with accuracy ranging from only 50.9 to 52.0%. However, the other four algorithms—QDA, RDA, MDA, and FDA—achieved high predictive accuracy in discriminating between BM and OU, with classification accuracies ranging from 95.3 to 98.1% across all conditions (Fig. 2). These methods were only minimally affected by measurement error in the trait data, consistently maintaining over 95% accuracy for both models. Additionally, all four algorithms produced balanced classification rates, showing no systematic bias toward either evolutionary model.

### Case Study II: Discriminating BM, OU, and EB

Our Case Study II analyses underscore the increased difficulty of selecting among three evolutionary models (BM, OU, and EB), compared to the two-model classification in Case Study I (Fig. 3). Predictive accuracy varied across algorithms and error conditions. Model selection accuracy of AIC was consistently lower than those of the top-performing EvoDA algorithms, ranging from 40.8 to 80.7% depending on conditions (Fig. 3). AIC accuracy was higher when error estimation was not included, with high accuracy for OU (99.5-99.8%), but substantially lower accuracy for BM (65.9-74%) and EB (16.2-68.7%). When error estimation was included, BM accuracy improved (76.5-76.8%), but at the expense of both OU (8.6-9%) and EB (37.1-67.1%). As with Case Study I, AIC (both with and without error estimation) was more sensitive to increasing measurement error in the trait data. Still, AIC outperformed LDA which appeared to choose classes with equal probability.

**FIGURE 3.**
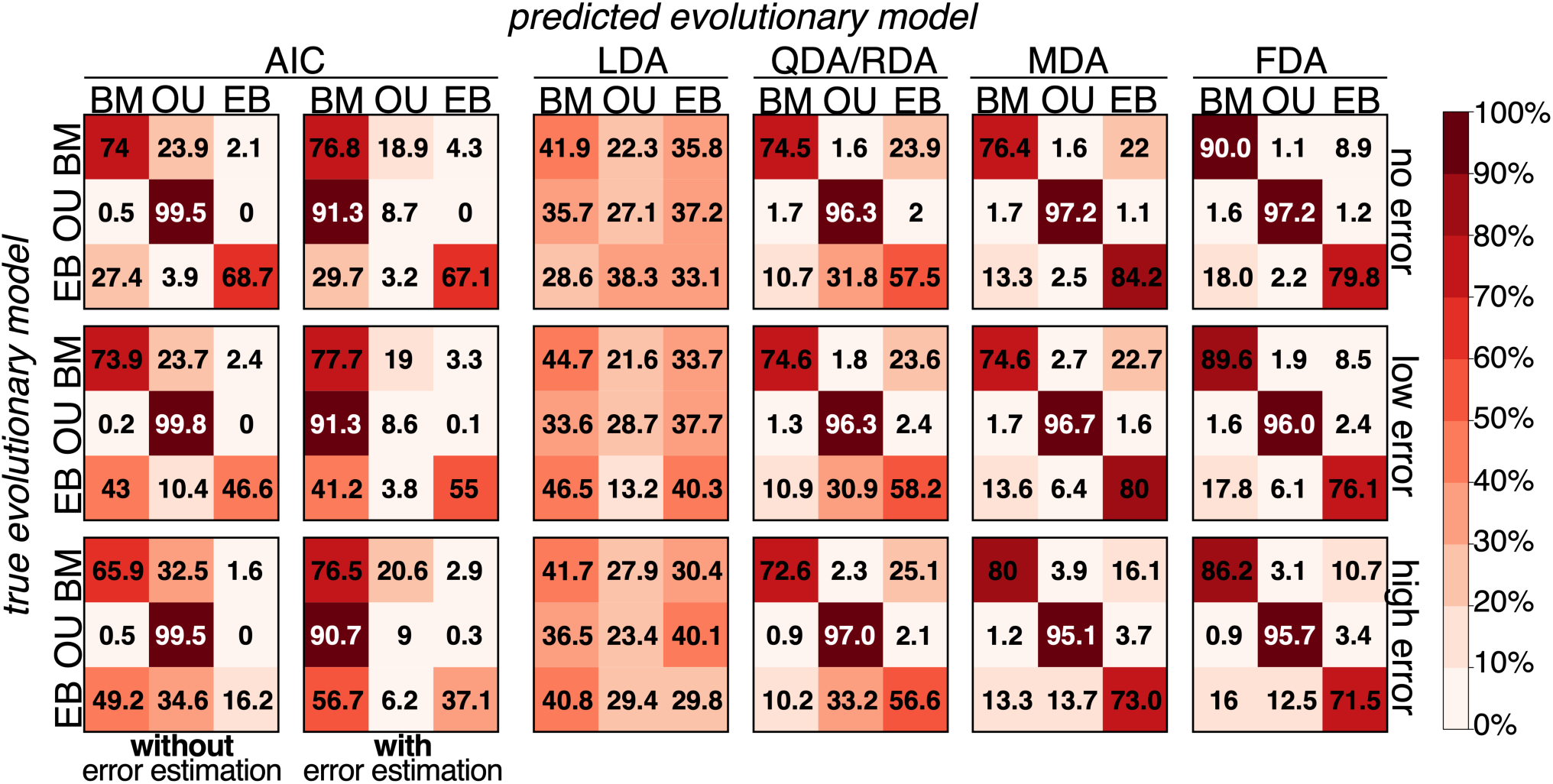
Case Study II: Discriminating among three canonical models of trait evolution. Confusion matrices (left to right) depict classification accuracies for AIC without and with error estimation, LDA, QDA, RDA, MDA, and FDA. QDA and RDA produced identical results and were therefore combined as QDA/RDA. Results are shown for experimental conditions with three levels of simulated trait measurement error: none (top row), low (middle row), and high (bottom row). In each matrix, rows represent the true evolutionary model used to simulate traits, and columns represent the predicted model. Shading intensity reflects the percentage of test replicates predicted for each model, with darker cells indicating higher values.

Across the EvoDA algorithms, FDA achieved the highest overall accuracy (84.5-89.0%), followed by MDA (83.0-86.0%), and QDA/RDA (75.4-76.1%). LDA again performed poorly with nearly equal probabilities of choosing BM, OU, or EB for any given trait. Accuracy was generally highest for OU (95.1-97.2%), moderate for BM (72.6-90.0%), and lowest for EB (56.6-84.2%). FDA outperformed MDA primarily by improving BM classification. Classification accuracy was slightly lower when analyzing data with measurement error (Fig. 3). For example, accuracy of FDA was 89.0% without error, which dropped to 84.5% under conditions with high error (Fig. 3), due to reduced accuracy for EB. Interestingly, MDA accuracy for BM slightly improved under conditions of high error.

### Case Study III: Discriminating seven canonical models

Case Study III revealed greater analytical challenges for all methods when selecting among a broader set of seven candidate evolutionary models (Fig. 4). Classification accuracy declined across all approaches in these conditions. When compared to the top-preforming EvoDA algorithms, AIC consistently yielded lower predictive accuracy across all conditions and error settings (16.4-35.3%). AIC performed best on the OU model when error was not estimated (76.4-78.8%), with the BM model recovering the second highest accuracy (54.8-63.6%) under these conditions. However, accuracy for the remaining models (EB, Kappa, Lambda Delta, and Trend) was generally quite low (0-29.4%). Including error estimation did not improve AIC performance; in many cases, trait models were misclassified as BM. Indeed, accuracy for many models remained below 20%, and many dropped to zero. As in our previous case studies, AIC also tended to be more sensitive to measurement error when compared with several of the EvoDA algorithms (Fig. 4).

**FIGURE 4.**
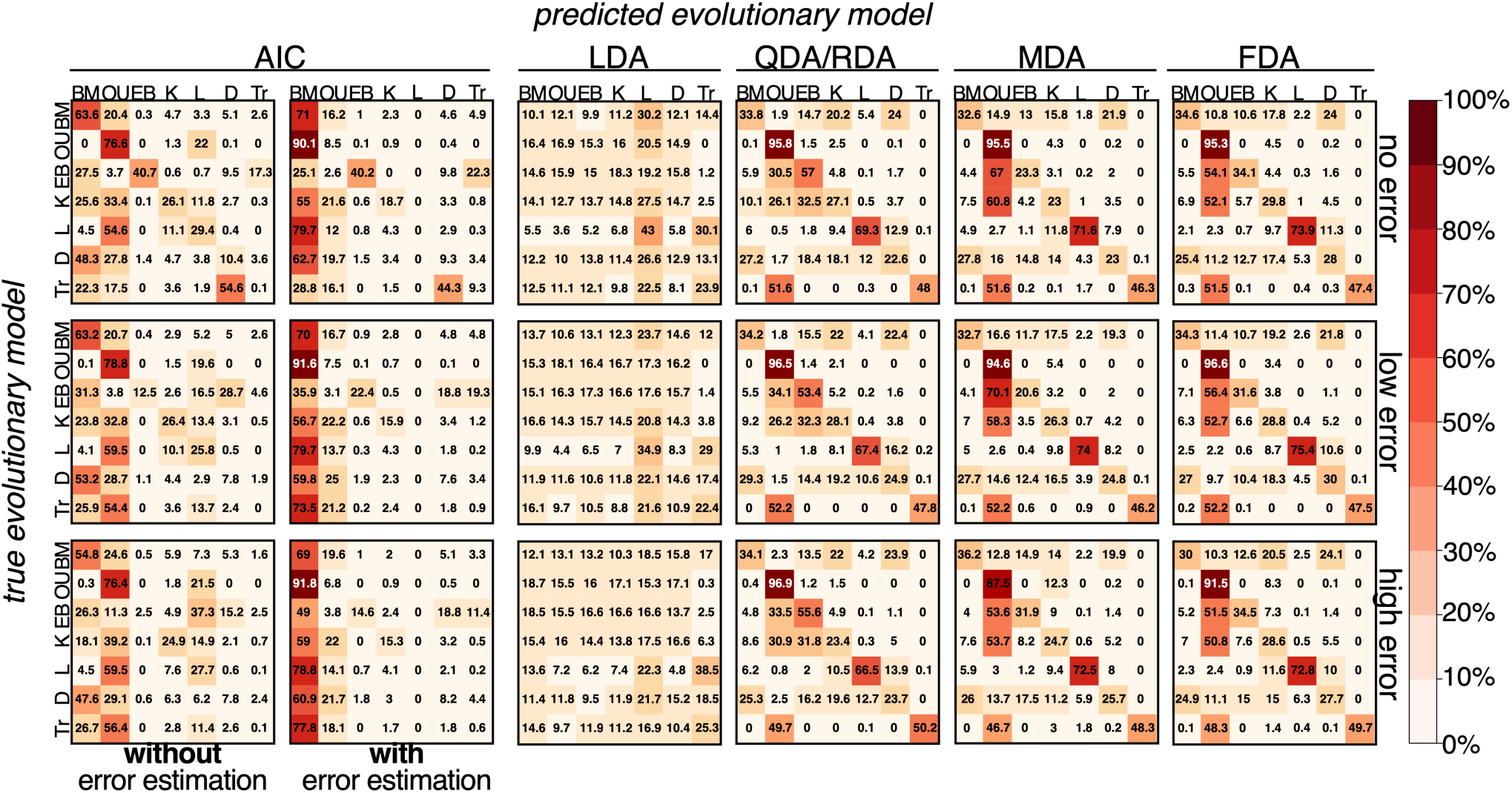
Case Study III: Discriminating among seven canonical models. Confusion matrices (left to right) depict classification accuracies for AIC without and with error estimation, LDA, QDA, RDA, MDA, and FDA. QDA and RDA produced identical results and were therefore combined as QDA/RDA. Results are shown for experimental conditions with three levels of simulated trait measurement error: none (top row), low (middle row), and high (bottom row). In each matrix, rows represent the true evolutionary model used to simulate traits, and columns represent the predicted model. Shading intensity reflects the percentage of test replicates predicted for each model, with darker cells indicating higher values.

Classification accuracy of EvoDA also declined, though four algorithms again achieved the highest accuracies: FDA (47.8-49.1%), MDA (45.6-46.7%), and QDA/RDA (∼50%). Among trait models, OU was consistently recovered with the highest accuracy (87.5-96.9%), followed by the Lambda model (66.5-75.4%). In contrast, EB (20.6-57%) and Trend (46.2-50.2%) showed more variable performance, while BM, Kappa, and Delta were predicted with substantially lower accuracy overall. Notably, EvoDA algorithms struggled to classify BM, reflected by low accuracies (30-36.2%), whereas QDA/RDA recovered the EB model with higher accuracy than either MDA or FDA. As in the previous case studies, predictive accuracies of EvoDA algorithms were not strongly influenced by measurement error, with only minimal reductions observed (Fig. 4).

### Empirical Case Study: learning gene expression evolution

Phenotypic variation among species may be shaped by differences in gene expression patterns and the evolutionary processes underlying them (King and Wilson 1975; Carroll 2005). The growing availability of multispecies gene expression datasets offers unprecedented opportunities to investigate these dynamics at scale (Chen et al. 2019). Yet, understanding the mode and tempo of expression evolution remains a persistent challenge (Cornwell and Nakagawa 2017). With this goal in mind, we applied and evaluated EvoDA for predicting the processes shaping the evolution of gene expression variation across fungal species using the same fungal phylogeny (Cope et al. 2020; Dimayacyac et al. 2023) employed in our simulation case studies. We also sourced the original RNA-seq data from these 18 fungal species from its published repository (Cope et al. 2020; Dimayacyac et al. 2023), which provided normalized, log-transformed expression counts for each species. Mirroring the design of Case Study II and consistent with previous fungal gene expression studies (Rohlfs et al. 2014; Chen et al. 2019; Cope et al. 2020; Dimayacyac et al. 2023), we focused on comparing the fit of three candidate models: BM, OU, and EB. To better align the training and testing data with the empirical expression data (see *Methods*), we employed the rejection sampling algorithm (see *Methods*; Campelo dos Santos et al. 2024).

Prior to applying the models to real gene expression data, we evaluated accuracy using test data subject to the same rejection sampling scheme as the training data (Fig. 5a). Under these conditions, the four top-performing EvoDA algorithms achieved high accuracies: FDA (90.3%), MDA (86.7%), and QDA/RDA (84.8%). FDA slightly outperformed MDA overall, primarily due to improved classification of BM (86.4% versus 73.5%), though it showed slightly lower accuracy for EB (88.1%) compared to MDA (90.5%). AIC performed less accurately overall (85.7%) than the EvoDA algorithms, again reflecting lower accuracy for BM (Fig. 5a). However, it achieved high accuracy for OU (99.7%), surpassing all other methods in that category. The performance gains for FDA largely stemmed from higher accuracy for BM (86.4%) compared to AIC (69.9%; Fig. 5a).

**FIGURE 5.**
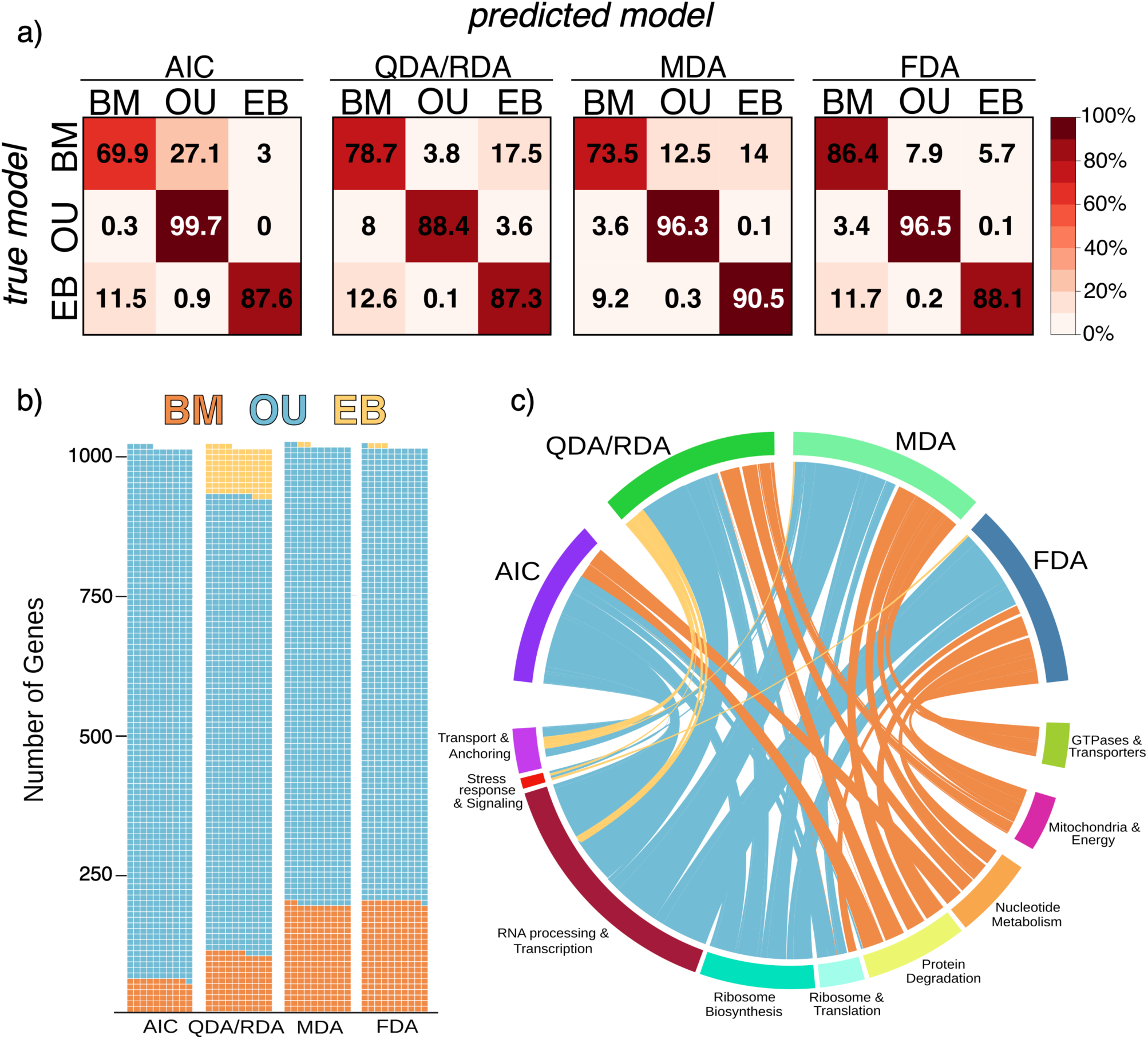
Learning gene expression evolution with EvoDA. (a) Confusion matrices depict classification accuracies of AIC, QDA/RDA, MDA, and FDA. In each matrix, rows represent the true evolutionary model used to simulate traits, and columns represent the predicted model. Shading intensity reflects the percentage of test replicates predicted for each model, with darker cells indicating higher values. (b) Predicted evolutionary models for 1,014 genes and (c) their associated biological function grouped by method and predicted model. Cell colors in (b) and connection bands in (c) correspond to the predicted model: BM (orange), OU (blue), and EB (yellow).

Our empirical application of EvoDA demonstrated strong potential for learning models of gene expression evolution. Consistent with previous studies (Cope et al. 2020; Dimayacyac et al. 2023), we found that a substantial proportion of fungal genes appear to evolve under OU processes, as reflected in model predictions across all focal algorithms (Fig. 5b). While OU was the most frequently selected model, we also observed evidence for BM and EB, with predictions varying by method. For example, AIC selected the BM model for 59 genes (5.8%), while QDA/RDA, MDA, and FDA predicted BM for 106 (10.45%), 192 (18.93%), and 199 (19.62%) genes, respectively. Though AIC did not select the EB model for any gene, QDA/RDA predicted EB for 87 genes (8.57%), while MDA and FDA predicted EB for two and three genes, respectively.

Genes and their predicted evolutionary models were linked to biological functions using Gene Ontology (GO) terms (Fig. 5c). Across the model selection procedures, the majority of genes were classified under the OU model, suggesting that stabilizing selection is the dominant evolutionary process shaping a wide array of cellular functions in fungi (Fig 5c). In contrast, genes assigned to the BM model were associated with a more restricted set of functions, including protein degradation, nucleotide metabolism, mitochondrial activity, and GTPase-mediated transport. Only a few genes were classified under the EB model, with functional annotations related to transport and anchoring, stress response and signaling, and RNA processing and transcription.

## Discussion

Evolutionary inference is inherently a retrospective task that seeks to reconstruct the past by linking observed trait variation with unobserved processes and patterns of trait change. For the vast majority of traits, it is impossible to know the precise evolutionary processes shaping variation across species. Instead, we rely on our ability to robustly and rigorously evaluate competing hypotheses to explain present-day trait variation in light of plausible past evolutionary scenarios. Guided by this goal, our study introduced EvoDA, a suite of supervised learning algorithms for predicting models of trait evolution through discriminant analysis, offering a new avenue for evolutionary inference. We view EvoDA as complementary to classical PCMs, with clear improvements in accurately predicting trait models in the presence of measurement error, reflecting realistic conditions in empirical studies. More broadly, our study also highlights persistent analytical challenges of inferring the mode and tempo of evolutionary processes, and the need for new strategies for learning trait evolution.

Perhaps the most evident advantage of EvoDA lies in its robustness to measurement error compared to conventional PCM-based model selection. Our findings align with previous studies demonstrating the sensitivity of PCMs to noisy data (Arnold 2010; Chakrabarti and Ghosh 2011; Brewer et al. 2016; Susko and Roger 2020; Bartoszek et al. 2023). By incorporating measurement error directly into the training process, we find that EvoDA is equipped to handle error while finding decision boundaries despite added noise, which likely reflects realistic empirical conditions. Measurement error is indeed a widespread concern for comparative studies (Ives et al. 2007; Villemereuil et al. 2012; Silvestro et al. 2015) because, at a fundamental level, biological variables are always measured with some degree of imprecision (Martins and Hansen 1997; Ives et al. 2007; Silvestro et al. 2015; Adams and Collyer 2018). In contrast, conventional model selection tends to be more sensitive to error, and we found that the option to estimate error did not readily rescue the analyses. Generally, AIC with measurement error tended to be biased toward the simple BM model, which was a pattern shared across the three case studies. Additionally, it is worth noting that most models of trait evolution are formulated to describe the evolution of the mean trait value of an entire species (Kelly and Price 2004; Ives et al. 2007). Yet, PCMs are often applied to traits measured from only a single individual or a small subset of individuals per species. As a result, some error is likely to be imposed by modeling the mean of a species based on few representative samples (Kelly and Price 2004; Ives et al. 2007).

The performance of the five EvoDA algorithms varied depending on the experimental conditions and evolutionary models considered. Across our study, both FDA and MDA tended to outperform other approaches, followed by QDA/RDA, and lastly, LDA. Clearly, LDA cannot be recommended, as this algorithm suffered from low accuracy, likely resulting from its overly simplistic assumption of equal covariance shared across classes (Park and Park 2008; Tharwat et al. 2017). Often, LDA appeared to simply choose classes at random across our simulation conditions. The clear improvements of QDA over LDA likely stem from increased flexibility of QDA through the incorporation of class-specific covariance structures, rather than a shared structure across classes. Such improvements of QDA have been seen in other applications (Ghosh et al. 2021). Throughout our analyses, RDA converged to QDA with proportion *α* = 1 of estimated covariance structure stemming from QDA, providing identical predictions and accuracy recovered from the two algorithms. This finding suggests that regularization with RDA favored the more complex QDA when selecting among evolutionary models, which is unsurprising as the covariance structure for all the trait evolution models considered differ. By incorporating additional complexity (Hastie et al. 1994), MDA often improved upon QDA/RDA by considering within-class variability, which is reflected in our broad range of parameter space (Table 1). The high preforming FDA likely benefited from the increased flexibility of nonlinear basis expansions that can capture more complex class boundaries (Bashir and Carter 2005; Reynès et al. 2006; Gkalelis et al. 2011). These findings underscore the importance of selecting discriminant methods that align with the complexity of the model selection task itself.

**TABLE 1.** Trait models, parameters, evolutionary processes, and sampling distributions used for training and test replicates. Distributions are based on the default parameter bounds assumed by the *fitContinuous* function in geiger.

| Model | Parameters | Evolutionary process | Distributions |
| --- | --- | --- | --- |
| BM | $z_0; \sigma^2$ | Random-walk model of trait change (Felsenstein 1973). | $\sigma^2 \sim \text{Exp}(1)$<br>$z_0 \sim N(0, 1)$ |
| OU | $z_0; \sigma^2; \alpha$ | Stabilizing selection (Butler and King 2004). | $\sigma^2 \sim \text{Exp}(1)$<br>$z_0 \sim N(0, 1)$<br>$\alpha \sim U(\exp(-500), \exp(1))$ |
| EB | $z_0; \sigma^2; a$ | Adaptive radiation: rates decline over time (Harmon et al. 2010). | $\sigma^2 \sim \text{Exp}(1)$<br>$z_0 \sim N(0, 1)$<br>$a \sim U(-5/\text{depth}, 10^{-6})$ |
| Lambda | $z_0; \sigma^2; \lambda$ | Phylogenetic signal scaling (Pagel 1999). | $\sigma^2 \sim \text{Exp}(1)$<br>$z_0 \sim N(0, 1)$<br>$\lambda \sim U(\exp(-500), 1)$ |
| Delta | $z_0; \sigma^2; \delta$ | Time-dependent evolutionary rate (Pagel 1999). | $\sigma^2 \sim \text{Exp}(1)$<br>$z_0 \sim N(0, 1)$<br>$\delta \sim U(\exp(-500), 3)$ |
| Kappa | $z_0; \sigma^2; \kappa$ | Branch length scaling in phylogeny (Pagel 1999). | $\sigma^2 \sim \text{Exp}(1)$<br>$z_0 \sim N(0, 1)$<br>$\kappa \sim U(\exp(-500), 1)$ |
| Trend | $z_0; \sigma^2; \text{slope}$ | Linear change in trait evolution over time (Pennell et al. 2014). | $\sigma^2 \sim \text{Exp}(1)$<br>$z_0 \sim N(0, 1)$<br>$\text{slope} \sim U(-100, 100)$ |

**TABLE 2.**
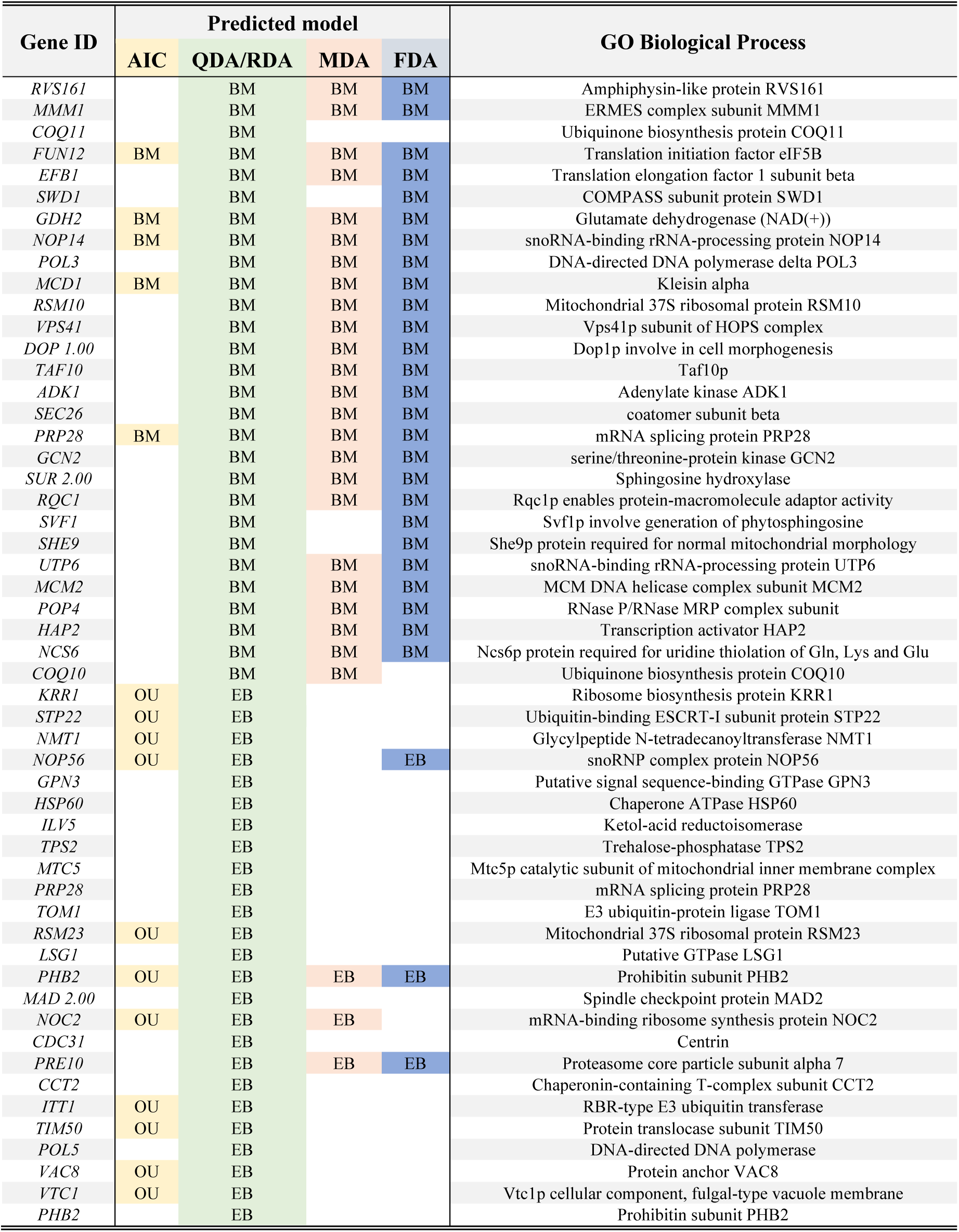
Evolutionary model predictions for genes with conflicting results across the four methods (AIC, QDA/RDA, MDA, and FDA), along with their associated GO Biological Processes.

Our study carries broader implications by highlighting the analytical challenges involved in selecting models of trait evolution, regardless of the technique employed. Clearly, the difficulty of model selection is likely influenced by many factors, including the space of candidate models considered, the presence and degree of measurement error, and for AIC, the option to estimate error or not. Importantly, all methods explored here struggled to perform well as the space of candidate models expanded. For example, four EvoDA algorithms (FDA, MDA, QDA, and RDA) performed exceptionally well at distinguishing BM versus OU, even in the presence of highly inaccurate trait measurements. Yet, classification accuracy dropped for all algorithms (and AIC) if only a single additional EB model was included in the candidate set, with even further reductions in accuracy when seven models were considered. Though we focused on canonical models that are commonly used in trait studies, alternative models have been proposed in the last decade (e.g., Landis et al. 2013; Rohlfs et al. 2014; Martin and Lenormand 2015), representing worthwhile considerations for future investigations. For tractability and consistency, we also focused our empirical analysis on a single, well-studied phylogeny of 18 fungal species. Future studies are needed to evaluate model selection techniques across a broader range of phylogenetic trees, taxonomic groups, and trait types.

Our empirical case study illustrates a promising application of EvoDA for learning mechanisms of gene expression evolution. Previous studies have lamented over the challenges of modeling the evolution of gene expression in fungi (Cope et al. 2020; Wint et al. 2022) and other systems (Romero et al. 2012; Munro et al. 2022). Our rejection sampling approach helped align EvoDA training and testing with properties of the empirical data, and such improvements with this strategy have been uncovered before (Campelo dos Santos *et al*. 2024). Under these conditions, several EvoDA algorithms recovered higher predictive accuracy compared to AIC-based model selection. While we focused on gene expression in this case study, traits vary widely in their statistical properties and genetic architectures. We therefore anticipate that rejection sampling may be broadly useful for tailoring simulations to empirical data across diverse trait types and research questions.

The evolution of gene expression is generally thought to reflect stabilizing selection that favors optimal expression values (Schraiber et al. 2013; Rohlfs et al. 2014), and previous studies have recovered this pattern, with many genes showing evidence of OU processes (Artieri and Fraser 2014; Dimayacyac et al. 2023). Comparative analyses focused on gene expression in yeast also indicate a widespread role of stabilizing selection on many fungal genes (Artieri and Fraser 2014). Notably, our empirical EvoDA applications also suggest a role for stabilizing selection on expression evolution governed by OU processes for a majority of genes (>80%). Yet, these analyses also revealed evidence of EB and BM for some genes (∼1-20%), which has been found in previous studies of fungi (Cope et al. 2020; Dimayacyac et al. 2023). Several EB-classified genes were associated with stress responses, aligning with previous studies suggesting that such genes tend to reflect rapid adaptive shifts (Romero et al. 2012). In contrast, BM-classified genes included those linked to protein degradation, nucleotide metabolism, mitochondria, and energy production, a pattern consistent with previous work suggesting that core metabolic processes often lack strong stabilizing pressure (Lynch et al. 2016).

Our investigation represents a series of case studies that focus on seven different algorithms for model selection, seven canonical trait models, a fungal phylogeny, three scenarios of measurement error, and an empirical analysis for the predicting evolution of 1,014 empirical expression traits. These decisions were based on both realism and tractability of our study to provide a first perspective on discriminant functions for classifying models of trait evolution. Expanding these conditions to include different trees, models, experimental conditions, and diverse trait-types (including multi-trait models) would likely provide an important avenue for future investigations, though it is worth emphasizing the computational demands required in expanding the scale and scope of simulation and empirical investigations.

## Methods

### Classifying Evolutionary Hypotheses

Though differing in implementation, EvoDA shares the same overarching goals as conventional PCMs: to predict a particular class that best describes the processes shaping trait evolution. From this perspective, we treat different evolutionary models as candidate classes that explain the observed trait variation across a set of target species. We denote a set of trait measurements taken across *n* species with the input vector

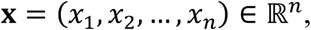

where *x_i_* is the value for the *i*th species. Due to shared evolutionary history, these trait measurements are not statistically independent (Freckleton et al. 2002). To address this issue, we transform the raw trait values into phylogenetic independent contrasts (PICs; Felsenstein 1985) redefining **x** as a vector of *m* = *n* − 1 contrasts:

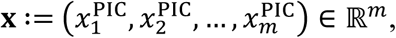

where 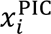 is *i*th PIC.

As with conventional model selection, we assume that the input vector **x** was generated under one of *K* possible classes, and we seek to predict its assignment to a class label **y** ∈ {1,2, …, *K*}. For example, we may wish to determine whether a trait evolved under a simple BM process (Cavalli-Sforza and Edwards 1967; Felsenstein 1973) or a more complex OU model (Lande 1979; Hansen 1997). For this study, we focus on selecting among seven canonical models of evolution: BM (Felsenstein 1973), OU (Hansen 1997), EB (Uyeda and Harmon 2014), Pagel’s Lambda, Delta, and Kappa (Pagel 1999), and rate trend (Trend; Mazel et al. 2016).

BM is the classical model of continuous trait evolution, which assumes that the mean trait value of a species evolves according to a single evolutionary rate parameter *σ*^2^ and an ancestral starting state *z*_0_ (Felsenstein 1973). The OU model extends BM to incorporate stabilizing selection by including an optimal trait value *z*) and a pull parameter *α* that quantifies the strength of attraction toward the optimum (Beaulieu et al. 2012; Tung Ho and Ané 2014; Cressler et al. 2015a). The EB model captures dynamics typical of adaptative radiations through a parameter *a*, which determines whether evolutionary rates increase (*a* > 0) or decrease (*a* < 0) exponentially over time (Harmon et al. 2010; Ingram et al. 2012; Slater and Pennell 2014; Martin et al. 2023). The Lambda, Delta, and Kappa models are formulated as tree transformations. The Lambda model modifies the tree topology using a parameter λ, scaling the tree from its original form (*λ* = 1) to a star phylogeny when (*λ* = 0). The Delta model alters the relative contributions of deep versus shallow nodes by raising node depths to the value of *δ*. The Kappa model adjusts all branch lengths by raising them to the power of the *κ* parameter. Finally, the Trend model extends BM by incorporating a linear trend in the mean trait value over time, allowing for directional evolution (Harmon et al. 2008). Collectively, these seven trait models have played prominent roles in comparative studies (Harmon et al. 2008; Eastman et al. 2011) and are widely applied across a variety of taxonomic systems (Henry et al. 2008), experimental conditions (Kawecki and Ebert 2004), and evolutionary questions (Pennell and Harmon 2013; Uyeda et al. 2018).

### Evolutionary Discriminant Analysis

The five EvoDA algorithms explored here include: LDA (Sudibyo et al. 2020; Zaki and Meira, Jr 2020), QDA (Finch and Schneider 2006; Sifaou et al. 2020; Ghosh et al. 2021), RDA (Zhu and Huang 2013; Yan et al. 2022), MDA (Reynès et al. 2006), and FDA (Ramsay and Silverman 2005). Each algorithm carries its own assumptions, strengths, and limitations (Tarca et al. 2007; Libbrecht and Noble 2015). These methods vary in their capacity to model deviations from multivariate normality and offer differing levels of flexibility, which are particularly relevant when working with a diversity of traits and phylogenetically transformed variables, as can be appreciated in our empirical case study (Fig. S1).

LDA is the most widely used discriminant function (Tharwat et al. 2017; Zhao et al. 2024), which can be defined as

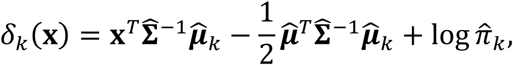

where 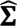 ∈ ℝ*^m^* is the estimated pooled covariance matrix that is shared across all *K* classes, 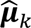 ∈ ℝ*^m^* is the estimated mean input vector for class *k*, 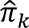 is the estimated prior probability of class *k*, superscript *T* indicates the transpose, and superscript −1 indicates the matrix inverse. LDA uses labeled training data to estimate the class-specific mean vectors 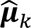, the shared covariance matrix 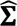, and the class priors 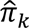. It assumes that class boundaries arise solely from differences in class means, while all classes share a common covariance structure. Under these assumptions, LDA finds linear decision boundaries that minimize overlap among class distributions centered on their respective means.

QDA extends the LDA framework by estimating a separate covariance matrix 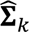 for each class *k* in addition to class-specific means 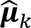 with the discriminant function

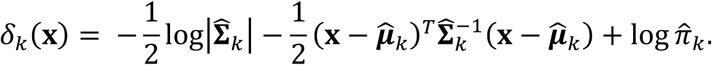

By relaxing the assumption of shared covariance across classes, QDA allows for greater flexibility in distinguishing among classes that differ not only in their means, but also in variance structure, distributional shape, and orientation in feature space (Ghosh et al. 2021). However, this flexibility comes at the cost of higher computational complexity and a greater risk of overfitting, particularly in small or high-dimensional datasets (Tarca et al. 2007; Cunningham et al. 2008). QDA is most effective when class distributions differ meaningfully in both their means and covariances, necessitating more complex, nonlinear (quadratic) decision boundaries (Wu and Hao 2022).

RDA introduces a regularization framework to strike a balance between LDA and QDA. Specifically, RDA computes a regularized class-specific covariance matrix as a weighted average of the LDA and QDA covariance estimates, controlled by a hyperparameter parameter *α* ∈ [0,1]:

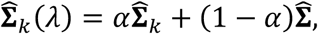

Where 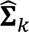 is the class-specific covariance matrix estimated in QDA, and 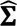 is the pooled covariance matrix used in LDA. RDA collapses to LDA when *α* = 0 and is equivalent to QDA when *α* = 1. RDA is particularly helpful in situations where the number of features exceeds the number of observations, or when the data exhibit high collinearity (Friedman 1989). By incorporating regularization into covariance estimation, RDA modulates the contributions of LDA and QDA, thereby balancing model complexity and providing more stable classification performance (Friedman 1989; Capblancq and Forester 2021). To determine the optimal value of *α*, we employed 10-fold cross-validation using an equally spaced grid search over *α* ∈ (0, 0.2, 0.4, 0.6, 0.8, 1), with highest classification accuracy as the selection criterion.

MDA extends the traditional discriminant analysis framework by allowing each class to be represented as a mixture of multivariate Gaussian distributions (Gkalelis et al. 2011). This approach helps capture intra-class variability by modeling each class as a combination of several subpopulations, each with its own mean and covariance matrix (Hastie et al. 1994; Tarca et al. 2007; Gkalelis et al. 2011). The probability density for an observation **x** given class **y** = *k* is defined by a Gaussian mixture model:

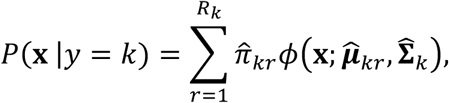

where *R_k_* is the number of mixture components for class *k*, 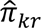 is the mixing proportion for the *r*th component in class *k*, and 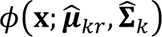 is the multivariate Gaussian density with mean 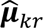 and class-specific covariance matrix 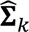. By incorporating within-class mixtures, MDA can model more complex class structures and overlapping subgroups, offering greater flexibility for challenging classification tasks. However, this increased complexity comes at the cost of higher computational demands and a greater risk of overfitting (Hastie et al. 1994; Gkalelis et al. 2011; Tharwat et al. 2017).

FDA employs nonlinear regression to define more flexible class boundaries (Hastie et al. 1994; Houdouin et al. 2023). Instead of relying on parametric assumptions, FDA employs nonparametric functions, such as B-splines (Reynès et al. 2006) and multivariate adaptive splines (Friedman 1991), to fit smooth, nonlinear decision boundaries. As a result, FDA can capture class boundaries that cannot be adequately represented by linear or quadratic forms (Hastie et al. 1994; Reynès et al. 2006), making it well-suited for classification tasks involving complex or irregular data structures (Hastie et al. 1994; Tharwat et al. 2017; Yang et al. 2018).

### Model Training and Validation

As a supervised learning framework, EvoDA algorithms are trained on labeled data in which input variables are matched with known output class labels representing evolutionary models. After training, predictive accuracy is assessed using “out-of-sample” test data that are not used during the training phase. Supervised learning is often a data-hungry process, requiring large datasets for both training and validation (Zhou 2018). To generate sufficient training and testing datasets, we leveraged the R package TraitTrainR (Roa Lozano et al. 2024), which simulates trait data under specified evolutionary models. For all EvoDA implementations, both training and test trait values were transformed into PICs prior to classification. In contrast, AIC-based model selection was conducted using standard approaches that incorporate phylogenetic information directly into the likelihood function. We also assessed two variants of AIC-based model selection that differed only in whether measurement error was estimated or not; we refer to these two settings as “AIC with error estimation” and “AIC without error estimation”, respectively. For our simulation case studies (Case Studies I-III; Fig. 1), we incorporated evolutionary variation throughout the algorithm training and testing procedures by randomly sampling parameter values for each model (BM, OU, EB, Kappa, Lambda, Delta, and Trend) across replicates (Table 1). Parameter bounds followed those used in the default implementations of the *FitContinuous* function from the geiger package (Pennell et al. 2014), which was also used for comparing models using AIC with and without measurement error. For our empirical case study, we also applied an additional rejection-sampling strategy (Hastie et al. 2009) to better align distributions of simulated training data with those of the observed gene expression data (see *Empirical Case Study* subsection below). For every analysis in our study, we simulated 10^5^ training replicates per class and 10^3^ out-of-sample test replicates per class to evaluate predictive accuracy.

### Predicting Trait Evolution in Three Simulation Case Studies

Case Study I evaluated the ability of EvoDA to discriminate between a simple BM process and a more complex OU model. Because there are only *K* = 2 classes, we refer to this case study as having an “easy” level of difficulty. Case Study II increased complexity by investigating the predictive performance of EvoDA for discriminating among *K* = 3 models: BM, OU, and EB, and we therefore refer to it as having a “medium” level of difficulty. Case Study III addressed the most complex scenario, requiring discrimination among *K* = 7 canonical models: BM, OU, EB, Kappa, Lambda, Delta, and Trend. These models vary in parameter count—BM has two parameters, while the others include three—and represent a diverse range of evolutionary processes and assumptions (Mazur 1959; Hansen 1997; Pagel 1999; Uyeda and Harmon 2014; Mazel et al. 2016). We thus refer to this case study as having a “hard” level of difficulty and use it to assess the performance of EvoDA when comparing multiple models with varying complexity, biological interpretation, and underlying dynamics (Arnold 2010; Brewer et al. 2016; Bartoszek et al. 2023).

### Empirical Case Study: Learning The Evolution of Gene Expression

To more closely align our training and testing simulations with the empirical expression data, we performed rejection sampling (Casella et al. 2004; Rubinstein and Kroese 2016) prior to EvoDA training and validation. Expression measurements for each gene were first transformed into PICs using the fungal phylogeny, yielding a vector of PIC values for each gene. As a filtering criterion, we used the standard deviation of PICs to exclude simulated trait replicates whose variance was abnormally high relative to the empirical data. Specifically, we computed the standard deviation of PICs for each of the 1,014 genes, resulting in a distribution with a mean of 0.0351 and a standard deviation of 0.0146. During the simulation process, we rejected simulated replicates that were outside the range of 0.0351 ± 3 × 0.0146 (mean ± 3 × standard deviation). This filtering ensured that retained simulations reflected the scale of empirical variation. From this procedure, we retained 10^5^ training replicates and 10^3^ testing replicates per class. Distributions of PICs for both empirical and filtered gene expression data are shown in Figure S1. Based on the poor accuracy of both LDA and AIC with error estimation in our simulation studies (see *Results*), we focused our empirical case study on the subset of the five best-performing methods: AIC without error estimation, QDA, RDA, MDA, and FDA. The four EvoDA algorithms were trained on the full set of retained training data and evaluated using the filtered test replicates to assess out-of-sample predictive accuracy. After rejection sampling, training, and testing, each model selection procedure was applied to the empirical data to predict the best evolutionary model for each gene.

## Funding And Acknowledgements

This research was supported by a grant from the Arkansas Bioscience Institutes, as well as the Arkansas High Performance Computing Center, which is funded through multiple National Science Foundation grants and the Arkansas Economic Development Commission. This work was also supported by National Science Foundation grant NSF DBI-2130666, National Institutes of Health grants R35GM142438 and R35GM128590, and start-up funds provided by the University of Arkansas.

## Resource availability

The software and framework EvoDA is available as an open-source R repository that includes all necessary tools for applying LDA, QDA, RDA, MDA, and FDA to trait evolution problems. The package includes comprehensive documentation and example datasets to facilitate use (Fig. 1). EvoDA builds on several widely used R libraries, including MASS (Venables and Ripley 2002), klaR (Weihs et al. 2005), mda (Hastie and Tibshirani 1998), and caret (Kuhn 2008). The package is freely available on GitHub at https://github.com/radamsRHA/EvoDA.

